# Destabilization of β-cell FIT2 by saturated fatty acids contribute to ER stress and diabetes

**DOI:** 10.1101/2021.02.28.433270

**Authors:** Xiaofeng Zheng, Qing Wei Calvin Ho, Minni Chua, Olga Stelmashenko, Sneha Muralidharan, Federico Torta, Elaine Guo Yan Chew, Michelle Mulan Lian, Jia Nee Foo, Markus Wenk, David L. Silver, Per-Olof Berggren, Yusuf Ali

## Abstract

Western type diets are linked to obesity and diabetes partly because of their high saturated fatty acid (SFA) content. We found that SFAs, but not unsaturated fatty acids (USFAs), reduced the number of lipid droplets (LDs) within pancreatic β-cells. Mechanistically, SFAs but not USFAs disabled LD biogenesis by inducing palmitoylation and subsequent ERAD-C mediated degradation of LD formation protein, Fat storage-Inducing Transmembrane protein 2 (FIT2). Targeted ablation of FIT2 reduced β-cell LD numbers, lowered β-cell ATP levels, reduced Ca2+ signaling, downregulated β-cell transcription factors (RNA sequencing analysis), and exacerbated diet-induced diabetes in mice. Subsequent mass spectrometry studies revealed increased C16:0 ceramide accumulation in islets of mice lacking β-cell FIT2 under lipotoxic conditions. Inhibition of ceramide synthases ameliorated the enhanced ER stress. Overexpression of FIT2 increased number of intracellular LDs and rescued SFA-induced ER-stress and apoptosis thereby highlighting the protective role of FIT2 and LDs against β-cell lipotoxicity and diet-induced diabetes.

## INTRODUCTION

Numerous evidences show that saturated fatty acids (SFAs) trigger ER stress, cell dysfunction and apoptosis in a myriad of cell types (1, 2). In contrast, unsaturated fatty acids (USFAs) remain relatively benign with some reporting opposite effects to SFAs (3, 4). The mechanism underlying this difference remains unclear. This is especially true for a cell-type as important as the pancreatic β-cell, where its vulnerability to fatty acid (FA) (i.e. lipotoxicity) reduces insulin provision and leads to diabetes (5). β-cells, as with all eukaryotic cells, possess the ability to generate intracellular lipid-containing organelles known as lipid droplets (LDs)(6).

LDs are highly conserved cytosolic organelles that originate from ER. They contain a hydrophobic core of *de novo* assembled neutral lipids and cholesterol esters, surrounded by a monolayer of phospholipids and cholesterol (7–9). An accumulating body of evidence suggests that the biological roles of LDs are broader than initially anticipated. Beyond mere lipid storage containers, LDs were shown to act as hubs for innate immune defence, modulators of nuclear function and sites for protein storage and trafficking (10–12). The number of LDs present in a cell can dictate its energy rate, inflammation state, resistance to pathogens, ER stress, and cell death. LD biogenesis is a complex process involving many different ER resident enzymes, including those that are involved in the synthesis of neutral lipids and cholesterol esters such as DGATs (13). However, LD formation is largely dependent on a highly conserved tripartite ER protein machinery comprising of seipins, perilipins, and FITs (14, 15). A modification in each of these proteins affect LD biogenesis from the ER (13).

Despite considerable attention and the link to many diseases, the contribution of LDs to β-cell dysfunction and diabetes is just beginning to emerge. LDs are present in β-cells and notable differences in LD numbers were observed between rodent and human β-cells, with the latter increasing in numbers per β-cell with age (16). Our underpinning observation of a significant change in LD number between murine β-cells exposed to SFAs with those exposed to USFAs, prompted a detailed mechanistic study on how SFAs may directly influence LD numbers in β-cells, and in turn how this regulation impacts β-cell function and whole-body glucose homeostasis. In this process, we uncovered a mechanism by which SFAs reduce LD numbers in β-cells with impact on β-cell function and survival, as well as overall glucose homeostasis.

Among the different proteins important for LD formation, we found that only the Fat storage-Inducing Transmembrane protein 2 (FIT2/FITM2), was rapidly downregulated with physiological levels of SFAs, but not with USFAs. FIT2 belongs to a unique family of evolutionarily conserved ER-resident, 6-transmembrane domain, protein with its only other member being FIT1/FITM1 (17, 18). FIT1 is primarily expressed in skeletal muscle, with lower levels found in heart, while FIT2 is ubiquitously expressed in tissues albeit at different levels (17). The highest levels of FIT2 were reported to be in both white and brown adipose tissue and the critical role of FIT2 for LD formation was shown in skeletal muscle, adipose tissue, and the small intestine (17, 19–21). FIT2, a critical protein for LD formation, has been shown to promote ER lipid coalescence and LD formation, a function that is likely linked to its catalytic lipid phosphatase activity (15, 18, 22, 23). Most recently, FIT2 was implicated in ER protein dysregulation, ER structure maintenance and the maladaptive ER response in both yeast and mammalian cells (23, 24). Despite this, the role and importance of FIT2 in β-cell LD formation, function and survival, especially during lipotoxicity, remain unknown.

Herein, we confirmed FIT2 presence in β-cells of mouse islets and we showed that SFA affects FIT2 levels post-translationally through palmitoylation, increased MARCH6 (E3 ligase) association, and proteasomal degradation in mouse insulinoma MIN6 cells. This loss, mimicked in a βFIT2KO mice (β-cell specific deletion of FIT2), resulted in reduced β-cell LD numbers and diet-induced diabetes with exacerbated glucose intolerance due to a reduced insulin secretory response. We found ceramide accumulation, a key driver of β-cell ER stress, in islets of βFIT2KO mice. Rescue of FIT2 in MIN6 cells, through overexpression, partially restored LD biogenesis, reduced ER stress and ameliorated SFA-mediated β-cell apoptosis (i.e., lipotoxicity). Beyond the conceptual advance of diet affecting β-cell function, this study highlights restoration of the β-cell LD formation as a potential way to preserve its function, especially in states of obesity.

## RESULTS

### Palmitate prevents β-cell lipid droplet accumulation through FIT2

To recapitulate β-cell vulnerability to SFAs, we first exposed MIN6 cells to physiologically relevant levels of either oleate (300 µM) or palmitate (300 µM). Exposure to oleate led to increased lipid droplets whereas exposure to palmitate did not (Fig 1A, B). We next determined the relative abundance of proteins critical for LD formation following either oleate or palmitate treatment. In MIN6 cells, only the FIT2 protein was downregulated by palmitate (16:0) and stearate (18:0), with levels lower compared to that of BSA-treated control cells. However, FIT2 was not downregulated by the chain-length equivalent USFA, palmitoleate (16:1) and oleate (18:1), respectively (Fig 1C, Supplementary Fig 1A-D). The loss of FIT2 corresponded with an increase in C/EBP Homologous Protein (CHOP) protein levels, a major downstream pro-apoptotic arm of the maladaptive endoplasmic reticulum (ER) stress response. In addition to MIN6 cells, FIT2 protein was also observed to be 3-fold lower in pancreatic islets from hyperphagic, obese, and diabetic, db/db mice compared to their non-diabetic littermates (where FIT2 mRNA co-localized with insulin-positive pancreatic β-cells) (Fig 1D, E, Supplementary Fig 1E). The reduced FIT2 protein abundance in db/db islets corroborates with prior evidence of reduced LD number in β-cells from diabetic mice (16). To confirm the role of FIT2 in reduced β-cell LD, we generated MIN6 cells lacking FIT2 (Fit2 shRNA, with 90% knockdown efficiency) (Supplementary Fig 1F). In parallel, mice bearing floxed (FL) *Fit2* alleles were bred with knockin mice bearing Cre recombinase inserted into the Ins2 locus (RIP-Cre), to achieve β-cell deletion of FIT2 (βFIT2KO) (Supplementary Fig 1G, H). Concern of RIP-Cre “leakiness” in certain regions of the brain, specifically the hypothalamus (25), was addressed by showing no change in *Fit2* mRNA expression in extra-pancreatic tissues (Supplementary Fig 2A), and by reporting no observable changes in IPGTT, except for a modest but significant increase in glucose excursion in RIP-Cre mice compared to wild-type mice, a well-documented RIP-Cre observation (26). There were no changes in ITT between the different mouse lines used (i.e. wild-type, RIP-Cre, floxed-Fit2 and βFIT2KO) (Supplementary Fig 2B - F). FIT2 downregulation significantly reduced LD number both in MIN6 cells (Fig 1F, G) and in βFIT2KO mouse islets, and the difference was exacerbated when βFIT2KO mice were placed on a moderate-SFA Western diet (West; a diet where saturated fat accounts for 40% of total energy) for 25 weeks (Fig 1H, I). Taken together, these results suggest LD accumulation, previously observed in human β-cells (16), as an added feature that distinguishes a β-cell response to either oleate or palmitate. The failure to accumulate LDs following palmitate exposure correlates with a loss of FIT2 protein in β-cells.

**Fig 1:**
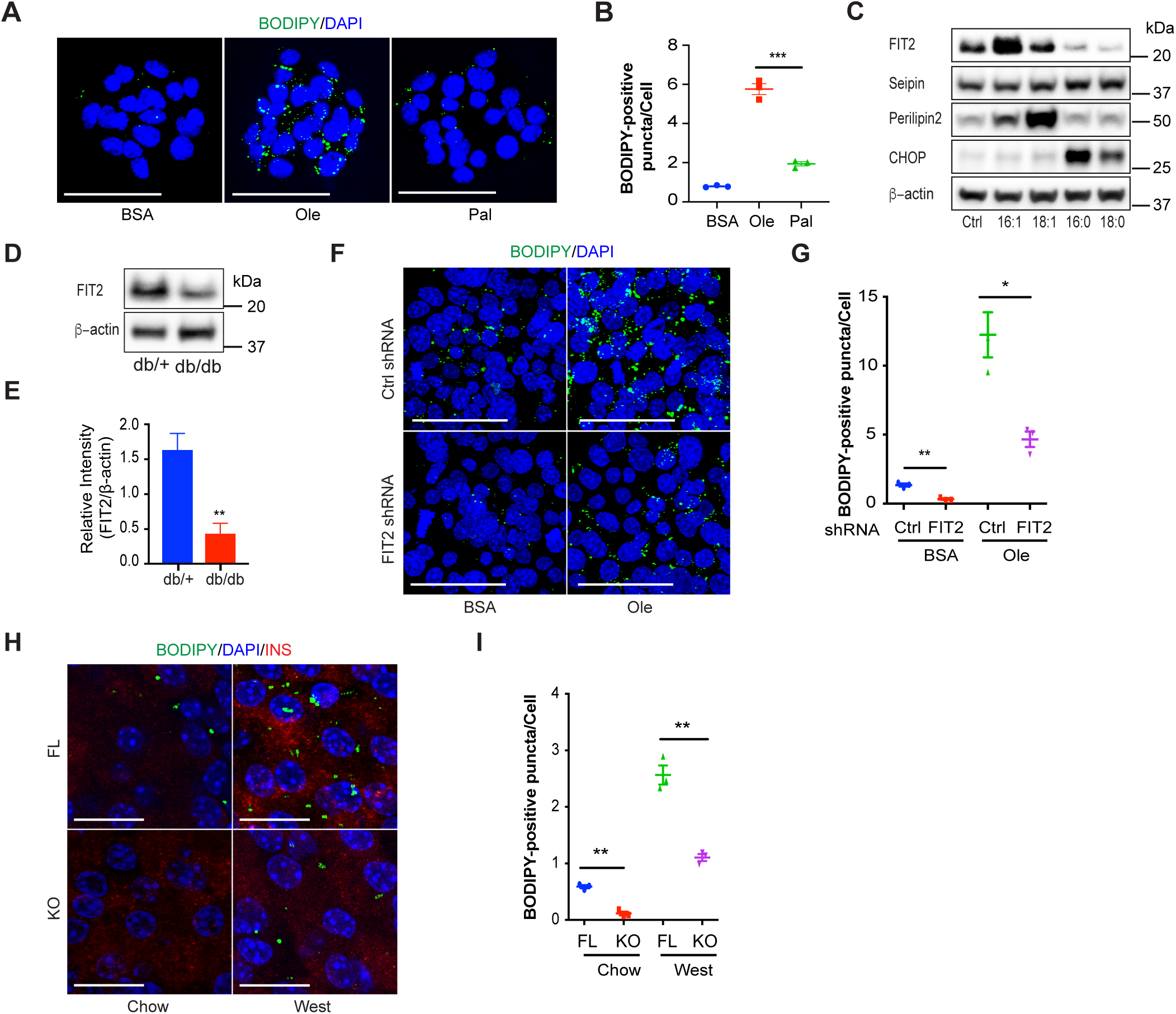
FA-mediated changes in β-cell LD numbers relates to FIT2 protein abundance. **A**, BODIPY staining of lipid droplets in MIN6 cells treated with BSA, oleate or palmitate (300 µM) for 24 h. Lipid droplets stained with BODIPY (green) and nuclei stained with DAPI (blue). Images shown are maximum-intensity projections. Scale bar = 50 µm. **B**, Quantitation of BODIPY-positive puncta per cell using Image-J. (>500 cells from 3 independent experiments were analyzed per group). **C**, Representative immunoblot of LD formation proteins after MIN6 cells treated with 300 µM of palmitoleate (16:1), oleate (18:1), palmitate (16:0), stearate (18:0) or BSA (control) for 24 h. **D**, **E**, Representative immunnoblot and semi-quantitation of FIT2 protein in islets isolated from db/+ and db/db mice (16 wk-old male, N=4). **F**, BODIPY staining of lipid droplets in control (Ctrl shRNA) or FIT2 knockdown (FIT2 shRNA) MIN6 cells treated with BSA or 300 µM of oleate for 48 h. Lipid droplets were stained with BODIPY (green) and nuclei were stained with DAPI (blue). Images shown are maximum-intensity projections. Scale bar = 50 µm. **G**, Quantitation of BODIPY-positive punctae per cell using Image-J. BSA-treated Control (Ctrl shRNA, blue) or FIT2 knockdown (FIT2 shRNA, red) and oleate-treated (300 µM) Control (Ctrl shRNA, green) or FIT2 knockdown (FIT2 shRNA, purple) (>1000 cells from 3 independent experiments were analyzed per group). **H**, BODIPY staining of lipid droplets in pancreas sections of chow fed floxed-control (FL, blue) and βFIT2KO (KO, red) and Western Diet (West, 25 wk) fed floxed-control (FL, green) and βFIT2KO (KO, purple). Lipid droplets were visualized using a BODIPY stain (green). β-cells were stained with insulin (red), and nuclei stained with DAPI (blue). Images shown are maximum-intensity projections. Scale bar = 25 µm. **I**, Quantitation of BODIPY-positive puncta per cell using Image-J (N=3, ≥9 islets from three mice were analyzed per group). Values shown are mean ± SEM; *, P < 0.05; **, P < 0.01; ***, P < 0.001 relative to control (two-tailed Student’s t-test).

### Loss of β-cell FIT2 exacerbates diet-induced diabetes

Since β-cell FIT2 levels were observed to be linked to FA exposure, we determined whether its loss impacted whole body glucose homeostasis. Loss of FIT2 in β-cells resulted in a modest but significant increase in plasma glucose concentration following bolus intra-peritoneal injection of glucose in chow-fed βFIT2KO mice (Fig 2A, B). However, the increased plasma glucose concentration observed in βFIT2KO mice was exacerbated in West diet fed (25 weeks) mice (Fig 2A, B). These results suggest that βFIT2KO mice were metabolically worse off when placed on a West diet as compared to West-fed floxed-control mice. Subsequent analysis showed that the poor glycemic control in βFIT2KO mice was driven by a significantly muted insulin response to high glucose, both *in vivo* and *ex vivo*, and not by changes in peripheral insulin sensitivity (Fig 2C – 2G). There were no discernible differences in islet morphology, islet cell composition, β-cell size, islet insulin content and insulin granule morphology between βFIT2KO and floxed control mice (Supplementary Fig 3A - F). Rather, the muted insulin response in βFIT2KO islets could be linked to an observed reduction in cellular ATP and a blunted glucose-stimulated intracellular Ca^2+^ influx (Supplementary Fig 3G – I). A subsequent comparative RNA-seq analysis between islet cells from βFIT2KO and floxed control mice revealed distinct gene expression changes in (i) vesicle fusion and exocytosis, (ii) ATP binding, (iii) Ca^2+^ transport and, (iv) ER-related genes as well as a significant downregulation of genes (such as Pdx1, MafA, Nkx6.1, Pax6) required for β-cell function (Fig 2H, I). These results lend support to earlier phenotypic observations of poor glucose control due to reduced β-cell function in βFIT2KO mice.

**Fig 2:**
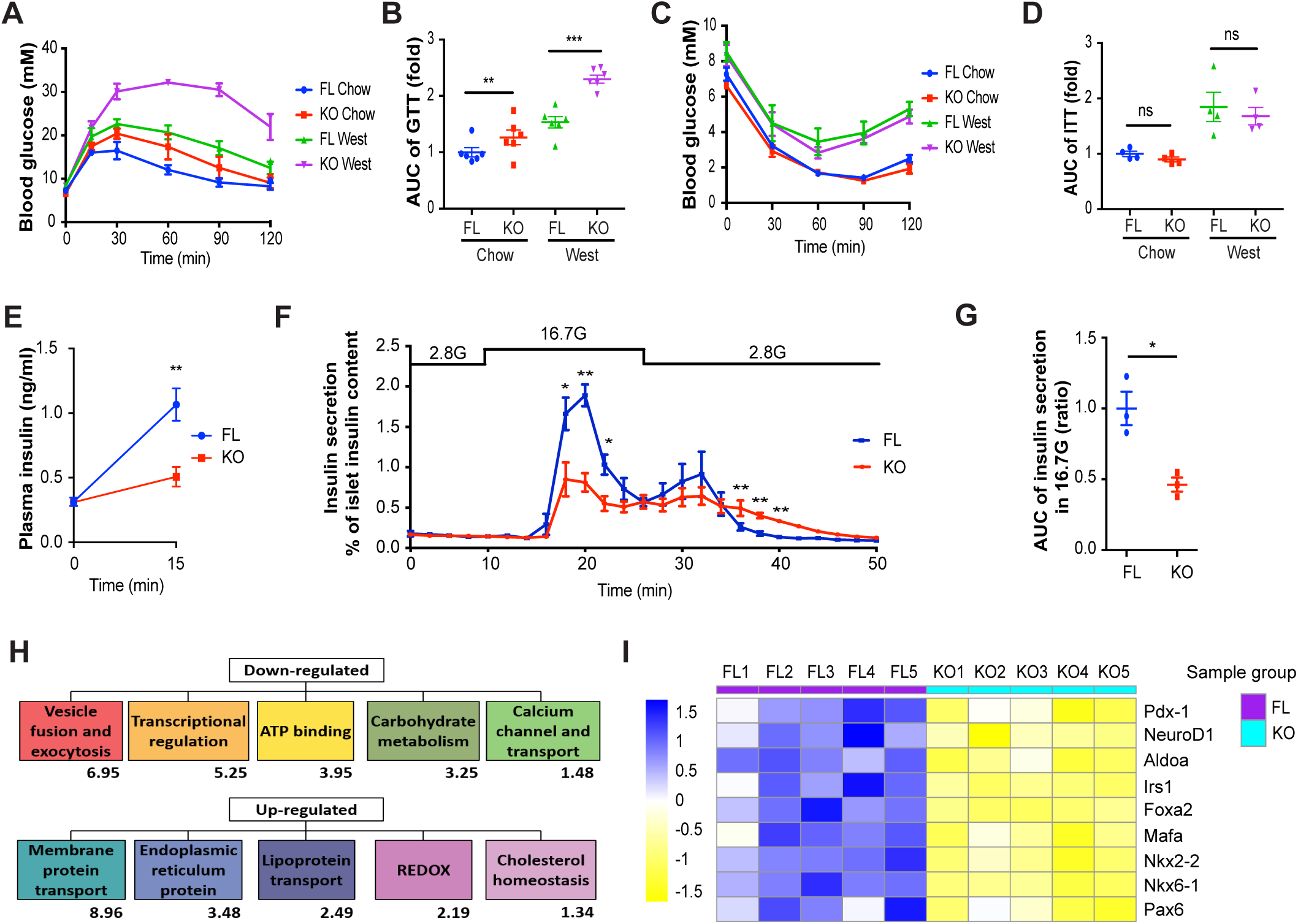
Glucose homeostasis and β-cell function is impaired in βFIT2KO mice. **A**, IPGTT on chow fed floxed-control (FL, blue) and βFIT2KO (KO, red) or Western Diet (West, 25 wk) fed floxed-control (FL, green) and βFIT2KO (KO, purple) and **B**, AUC of blood glucose levels from A (N=6). **C,** ITT on chow fed floxed-control (FL, blue) and βFIT2KO (KO, red) or Western Diet (West) fed floxed-control (FL, green) and βFIT2KO (KO, purple) **D,** AUC of blood glucose levels from C (N=4). **E**, *In vivo* insulin secretion following glucose administration (I.P.) in floxed-control (FL) and βFIT2KO (KO) mice (12 wk-old male, N=7). **F**, *Ex vivo* islet glucose-stimulated insulin secretion from floxed-control (FL) and βFIT2KO (KO) mice (12 wk-old male, N=3). **G**, Area under the curve (AUC) for insulin secretion at 16.7 mM glucose (16.7 G) from F (N=3). **H**, Gene ontology (GO) clustering analysis of down- and up-regulated genes in (βFIT2KO) KO as compared to floxed-control (FL) islets (N=4). Enrichment score of each GO cluster indicated below respective term, significant terms represented by enrichment score ≥ 1.3 (Supplementary Table 1). **I,** Expression of key β-cell genes in islets from FL and KO mice extracted from RNA-Seq data in H. Values shown are mean ± SEM; *, P < 0.05; **, P < 0.01; ***, P < 0.001 relative to control (two-tailed Student’s t-test).

### Loss of FIT2 increases β-cell ceramides and exacerbates ER stress

The inability of β-FIT2KO mice to cope with a chronic 25-week exposure to West diet was further scrutinized. We observed higher levels of p-IRE1α^+^/ insulin^+^ cells and ATF4^+^/ insulin^+^ cells in West-fed β-FIT2KO (KO West) mice compared to the West-fed floxed control (FL West) mice (Fig 3A - D). This suggests that loss of FIT2 in β-cells increased the maladaptive ER stress response *in vivo*. Exposure to FA (especially palmitate) triggers an ER stress response, an important driver of β-cell dysfunction (27, 28). Therefore, we determined whether β-cell FIT2 and LDs played any part in the palmitate-induced ER stress response. In FIT2 knockdown MIN6 cells (FIT2 shRNA), p-IRE1α was found to be significantly elevated, with a time-dependent increase in IRE-1, ATF4 and CHOP protein levels compared to control MIN6 cells (Fig 3E; Supplementary Fig 4A - D). The amplified ER stress response (ATF4, CHOP and IRE-1 pathways) coincided with increased cell death (Fig 3F, G), and a significantly elevated activity of caspase-3/7, suggesting increased apoptosis in palmitate-treated FIT2 shRNA cells compared to palmitate-treated control cells (Fig 3H). Liquid chromatography-mass spectrometry (LC-MS) analysis of ceramide levels in pancreatic islets isolated from KO West and FL West (fed) mice revealed accumulation of toxic C16 ceramide (Cer d18:1/16:0), in the β-FIT2KO islets (Fig 3I). Similarly, there was a modest increase in the ceramide precursor C16 dihydroceramide (Cer d18:0/16:0) (p=0.055) and in other dihydroceramides containing saturated fatty acyl chains, suggesting enhanced *de novo* ceramide synthesis in islets lacking FIT2. A toxic accumulation of ceramides in β-cells was corroborated by immunofluorescence (IF) analysis in β-FIT2KO mice (Supplementary Fig 4E and F) and co-treatment of palmitate with Fumonisin B1 (FB1), a pharmacological inhibitor of ceramide synthases, reduced CHOP levels (Fig 3J, K) (29). Together, these data suggest that ceramide accumulation contributed to enhanced ER stress and dysfunction in West-fed β-FIT2KO mice.

**Fig 3:**
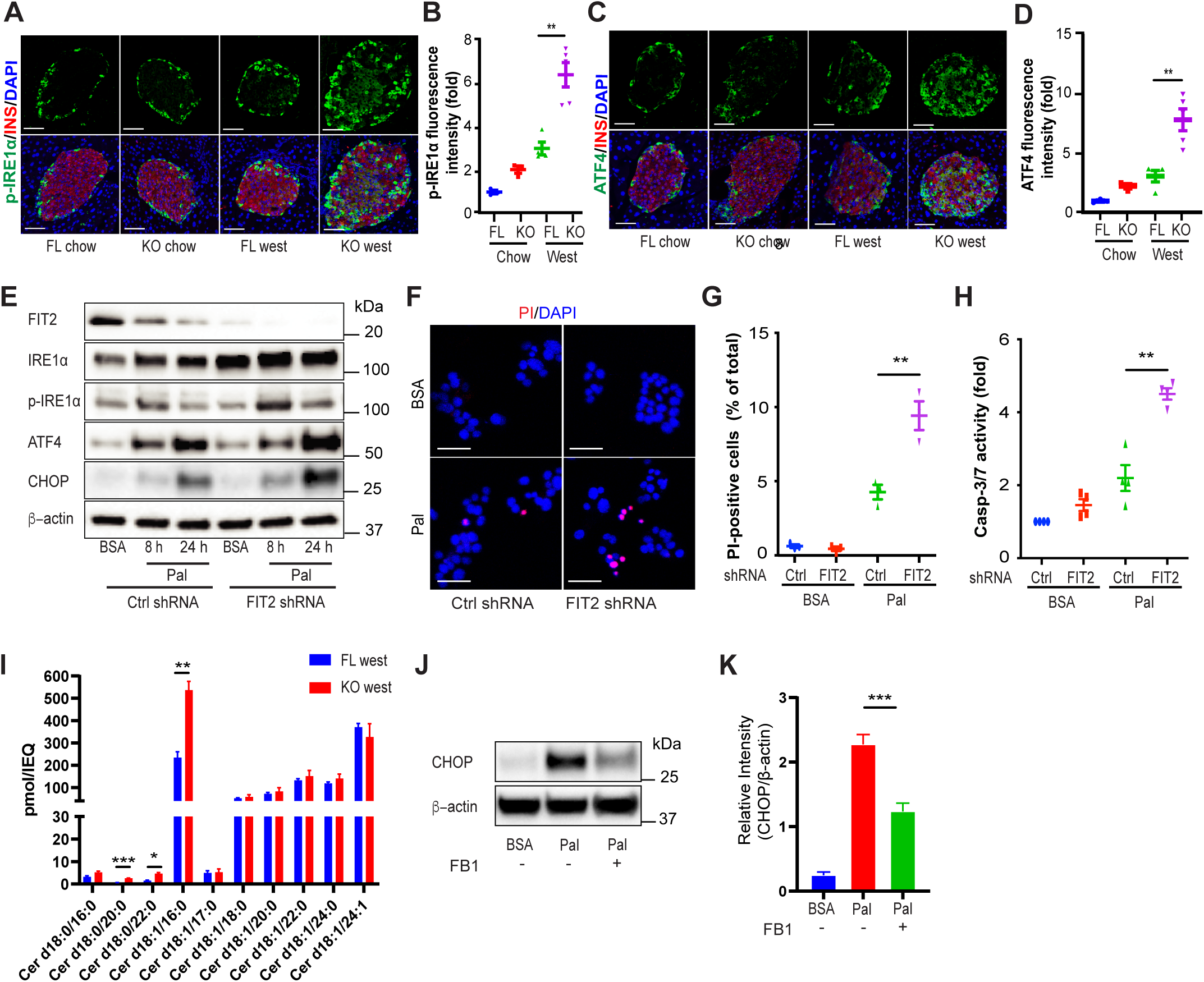
Loss of β-cell FIT2 exacerbates ER stress and survival. **A**, **C**, Representative immunostain for p-IRE1α and ATF4 (green), insulin (red) and DAPI (blue) in pancreas sections from floxed control (FL) and βFIT2KO (KO) mice treated with either Chow or West diet (25 wk). Images shown are maximum-intensity projections with a scale bar of 50 µm. **B, D**, Quantitation of relative fluorescence intensity of p-IRE1α and ATF4 staining within insulin-positive cells (N=3-5 mice, ≥9 islets from each mouse were analyzed per group). **E**, Representative immunoblot of indicated proteins in control shRNA transfected or FIT2 stable knockdown MIN6 cells (FIT2 shRNA) in the absence or presence of palmitate (300 µM). **F**, Representative image of control shRNA transfected or FIT2 stable knockdown (FIT2 shRNA) MIN6 cells in the absence (BSA only) or presence of palmitate (300 µM, 24 h) stained for DAPI (blue) and Propidium Iodide (PI, red). Images shown are maximum-intensity projections. Scale bar = 50 µm. **G**, Quantitation of the percentage of PI-positive cells over total number of cells (>1000 cells from 3 independent experiments were analyzed per group). **H,** Measurements of capase-3/7 activity in control shRNA transfected or FIT2 knockdown (FIT2 shRNA) MIN6 cells in the absence (BSA only) or presence of palmitate (300 µM, 24 h) (N=4). **I**, Quantitation of different ceramide species in islets from 25 wk of west-diet fed floxed mice (FL-west) and βFIT2KO (KO West) by LC-MS (N=3). **J, K,** Representative immunoblot and analysis of CHOP in MIN6 cells treated with Fumonisin B1 (FB1) (10 µM) in the presence and absence of palmitate (300 µM, 4h) (N=4). Values shown are mean ± SEM; ns, not significant; **, P < 0.01; ***, P < 0.001 relative to control (two-tailed Student’s t-test).

### Partial restoration of FIT2 rescues LD biogenesis and mitigates palmitate-mediated effects in MIN6 cells

The role of FIT2 loss in SFA-induced β-cell apoptosis was further investigated through its overexpression in MIN6 cells. MIN6 cells were transiently transfected with either an expression vector encoding FIT2 (pcDNA3.1-FIT2) or with a control vector (pcDNA3.1-mock) prior to palmitate (or BSA) exposure. FIT2 levels increased by approximately 3-fold in FIT2-overexpressing (FIT2-OE) cells treated with BSA, compared to control (mock cells) (Fig 4A, B). While palmitate exposure resulted in an approximately 95% reduction of FIT2 in mock cells, this was partially attenuated in FIT-OE cells (approximately 55% reduction) (Fig 4A, B). This suggests that overexpression of FIT2 partially compensated for palmitate-induced reduction in FIT2 protein levels. This partial restoration of FIT2 in FIT-OE cells exposed to palmitate led to a modest but significant increase in the number of LDs (Fig 4C, D), suggesting a partial rescue of palmitate-induced LD loss in β-cells. This rescue followed with a corresponding reduction in CHOP protein levels and lowered caspase-3/7 activity in FIT2-OE cells exposed to palmitate (Fig 4E, F). Together, these results suggest that partial compensation of FIT2 loss ameliorates palmitate-induced lipotoxicity in β-cells.

**Fig 4:**
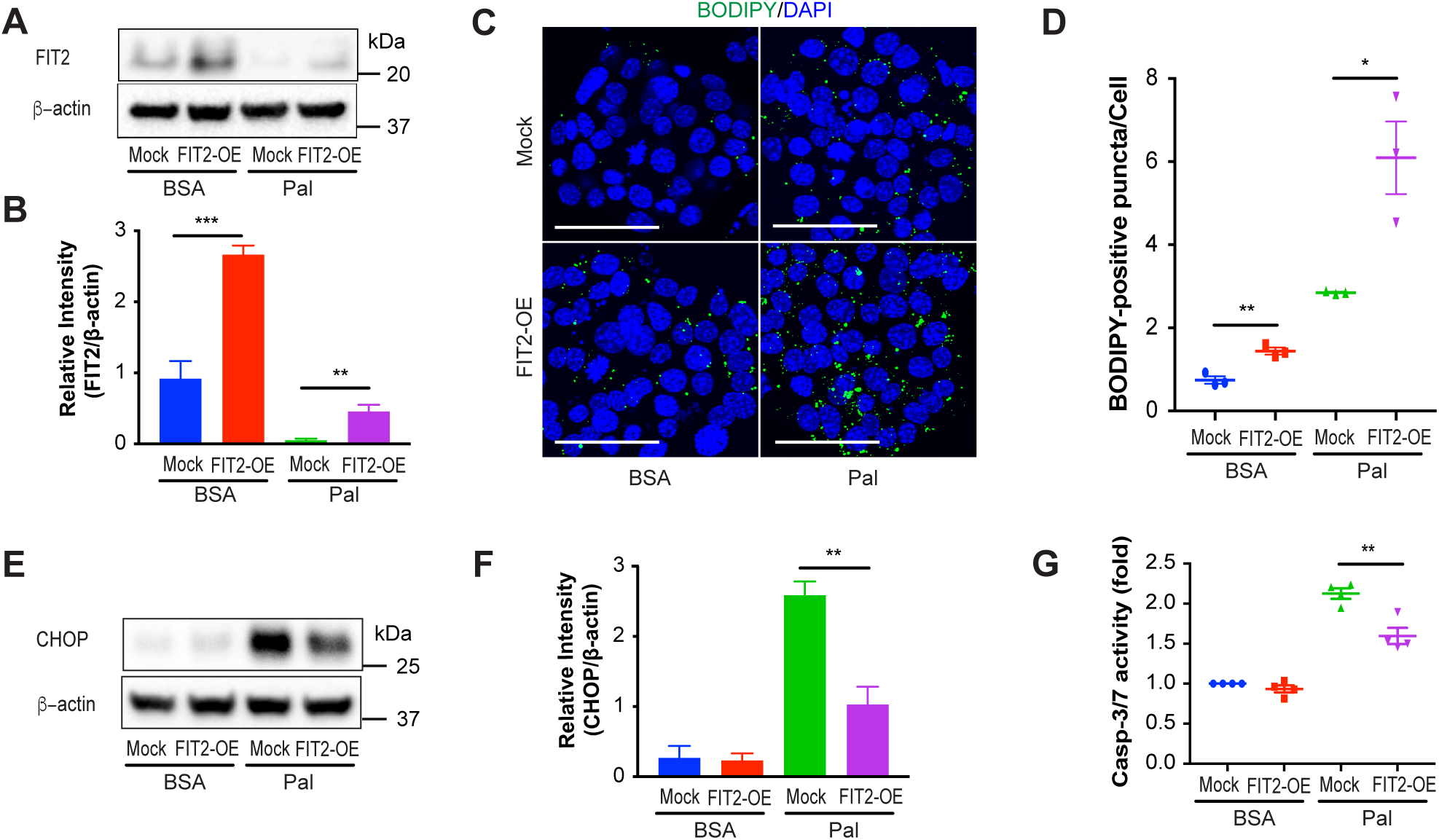
Overexpression of FIT2 in MIN6 cells ameliorates palmitate-induced lipotoxicity. **A-G**, MIN6 cells were cultured in 6-well plates and transfected with 0.5 µg empty pcDNA3.1 vector (Mock) or pcDNA3.1-FIT2 (FIT2-OE) per well, followed by treatment with BSA or palmitate (300 µM) for 24 h. **A**, **B**, Representative immunoblot and semi-quantitation FIT2 protein levels (N=4 independent experiments). **C**, Lipid droplet staining (BODIDY, green) in vector transfected (Mock) or in FIT2 overexpressing (FIT2-OE) MIN6 cells (nuclei stained with DAPI; blue) in the absence (BSA) and presence of palmitate (300 µM, 24 h). Images shown are maximum-intensity projections with a scale bar of 50 µm. **D**, Quantitation of BODIPY-positive puncta per cell using Image-J. (>1000 cells from 3 independent experiments were analyzed per group). **E**, Representative immunoblot analysis and **F**, corresponding semi-quantitation of CHOP levels (N=4). **G**, Measurements of capase-3/7 activity in control vector transfected (Mock) or FIT2 overexpressing (FIT2-OE) MIN6 cells in the absence (BSA) or presence of palmitate (300 µM) (N=4). Values shown are mean ± SEM; *, P < 0.05; **, P < 0.01; ***, P < 0.001 relative to control (two-tailed Student’s t-test).

### Palmitoylation and ERAD-C degradation pathways contribute to FIT2 protein loss

We next sought to elucidate the mechanism that may account for SFA-induced FIT2 loss. The observed SFA-mediated downregulation of FIT2 in MIN6 cells was not seen at the transcriptional level (Supplementary Fig 5A) but was abrogated in the presence of proteasome inhibitor MG132 (Fig 5A, B), suggesting that SFAs modulate FIT2 protein stability rather than its gene expression in β-cells. Given that FIT2 is an ER-resident protein, we then probed the possible involvement of an ER-associated degradation (ERAD) pathway in palmitate-mediated degradation of FIT2. ERAD inhibitor Eeyarestatin I (ES) (30), partially attenuated palmitate induced FIT2 loss (Fig 5C, D). Increased association between FIT2 and MARCH6, a mammalian E3 ligase complex responsible for degradation of the ERAD-cytosolic (ERAD-C) substrates (31), was observed in MIN6 cells treated with palmitate (Fig 5E). Furthermore, MARCH6 silencing (Supplementary Fig 5B) modestly but significantly rescued palmitate mediated FIT2 loss (Fig 5F, G) and taken together, these results implicate ERAD-C pathway and MARCH6 are involved in palmitate-mediated degradation of FIT2. The ERAD-C pathway is usually triggered by a change in the tertiary conformation of an ER-protein. We tested the possibility of palmitate directly modifying FIT2 given that an *in silico* analysis (GPS-lipid) (32) showed high probability of fatty-acid S-acylation taking place at 4 different cysteine residues (Cys-7, Cys-251, Cys-140 and Cys-70) (Supplementary Fig 5C). Indeed, the presence of S-acylation (palmitoylation) on FIT2 protein was confirmed with the S-palmitoylation assay, with higher levels of S-acylated FIT2 proteins detected in MIN6 cells exposed to palmitate (Fig 5H). Mutation of all 4 predicted S-acylation sites (Cys to Ala) abrogated palmitoylation under steady-state (BSA) and heightened (palmitate) conditions (Fig 5H). Next, to determine whether FIT2 palmitoylation led to its degradation, the palmitoylation inhibitor Cerulenin (33), was added to MIN6 cells together with palmitate. Cerulenin significantly restored palmitate-mediated FIT2 protein loss (Fig 5I, J). Given selectivity concerns of the palmitoylation inhibitor cerulenin, we further determined the stability of the FIT2 mutant (C7/251/140/70A) in the presence and absence of palmitate (Supplementary Figure 5C). Single mutation of either cysteine residue modestly, but insignificantly, abrogated palmitate induced FIT2 protein loss (Supplementary Fig 5D, E). However, mutation of all four cysteine residues completely blocked palmitate (Fig 5K, L) as well as stearate (Supplementary Fig 5F, G) mediated FIT2 degradation. Altogether, these results suggest that SFA-mediated S-acylation (palmitoylation) of the FIT2 protein promotes its degradation, in an ERAD-C and MARCH6 dependent manner. The loss of FIT2, in turn, triggers the maladaptive ER stress response and apoptosis in β-cells corroborating a previous finding in eukaryotic cells (23). This pathway that links SFA-mediated β-cell lipotoxicity with FIT2 degradation and reduced LD numbers has not been described previously.

**Fig 5:**
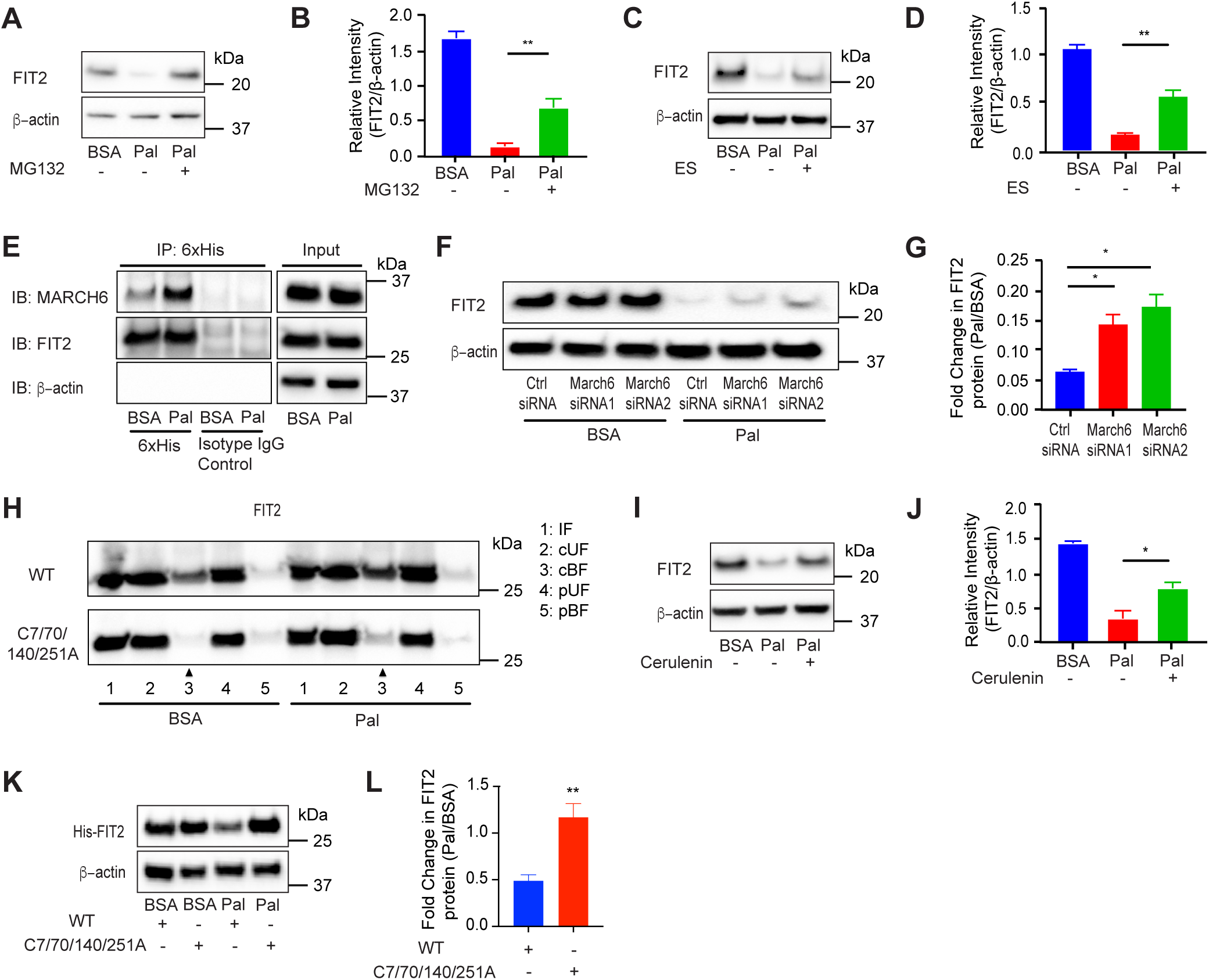
Palmitate-induced reduction of FIT2 protein involves palmitoylation and its ERAD-C mediated degradation in MIN6 cells. **A-D,** Representative immunoblot analysis and semi-quantitation of FIT2 protein in MIN6 cells pre-treated with respective pharmacological inhibitors (10 µM MG132 or 10 µM ES, 2 h), followed by co-treatment with inhibitors in the presence of palmitate (300 µM) for 4 h (N=3). E, Representative immunoblot for co-immunoprecipitation (Co-IP) of FIT2 and MARCH6 in MIN6 cells in the absence (BSA) and presence of palmitate (Pal, 300 µM, 6 h), following pre-treatment with 10 µM MG132 for 2 h. Pull-down of FIT2 using equal amount of input protein was carried out using 6xHis antibody (with isotype control) and thereafter captured proteins were probed for MARCH6 protein using the tGFP antibody. **F,** Representative immunoblot analysis and **G,** semi-quantitation of FIT2 protein from serum-starved MIN6 cells (2 h) transfected with scrambled siRNA (Ctrl siRNA) or March6 siRNAs, followed by treatment with BSA or palmitate (300 µM) for 4 h (N=4). **H,** Representative immunoblot of S-palmitoylation assay on both wildtype (WT) and mutant (C7/70/140/251A) FIT2 under steady-state (BSA) and palmitate (Pal, 300 µM, 6 h) conditions. The increased S-palmitoylation of FIT2 under palmitate conditions (lane 3, CAPTUREome™ resin cleaved bound fraction (cBF)) was markedly abrogated in the mutant FIT2. Input fraction (IF) and the flowthrough fraction which represents the cleaved unbound fraction (cUF) were collected and analysed. A separate set of protein lysates (Negative control) was treated with acyl-preservation reagent and subjected to incubation with CAPTUREome™ resin (Flowthrough fraction was used as the preserved unbound fraction (pUF) while eluted fraction was used as the preserved bound fraction (pBF)) **I,** Representative immunoblot and **J**, corresponding semi-quantitation of FIT2 protein levels in MIN6 cells pre-treated with Cerulenin (45 µM) for 2 h, followed by co-treatment of cerulenin (45 µM) in the presence of palmitate (300 µM) for 4 h (N=3). **K,** Representative immunoblot analysis and **L,** corresponding semi-quantitation of FIT2 protein levels in MIN6 cells transfected with pcDNA3.1-FIT2/V5-His (WT) or mutant (C7/70/140/251A) followed by treatment with BSA or palmitate (300 µM) for 24 h. Values shown are mean ± SEM; *, P < 0.05; **, P < 0.01; ***, P < 0.001 relative to control (two-tailed Student’s t-test).

## DISCUSSION

In the present study, we provide the first evidence of a FA-type specific modulation of ER resident, LD formation protein, FIT2 in β-cells. Our results show that LD biogenesis is affected by the type of FA exposure and that one pathway responsible for differential toxicity between dietary SFA and USFA (in the form of palmitate and oleate, respectively) involves cellular FIT2 protein stability, and the consequent ability of β-cells to accumulate LDs. We present a mechanism how SFAs reduce FIT2 levels leading to increased β-cell ER stress response, reduced insulin secretion and elevated β-cell apoptosis, as well as increased susceptibility to diet-induced diabetes. Altogether, we highlight the underlying novel finding that the ability of β-cells to withstand lipotoxicity is contingent on its capacity to maintain FIT2 and to generate intracellular LDs.

LDs are an important source for organelle membrane lipids and proteins (34). Mitochondrial lipid uptake from ER is essential for maintaining mitochondrial membrane integrity which may be affected by the loss of FIT2 as seen with decreased [ATP]/ [ADP] ratios (in βFIT2KO islet cells) (35). We observed significant elevation of ER stress-related proteins such as p-IRE1α and ATF4 in FIT2KD cells and islets of βFIT2KO mice. Corroborating transcriptomic evidence of ER protein and REDOX gene upregulation in βFIT2KO islets suggest that loss of LDs induces the maladaptive ER stress response in β-cells. Accumulation of C16:0 ceramides in islets of β-FIT2KO mice can be a strong contributor to the observed ER stress and β-cell dysfunction. A GENEVESTIGATOR® search revealed CerS5 and CerS6 to be the most abundant ceramide synthases in pancreatic islets (36). CerS5 and CerS6 generate C16 and C14 chain length ceramides, and are likely responsible for the observed difference in C16:0 ceramide levels in West-fed β-FIT2KO islets and West-fed floxed islets (37, 38). Further inhibition of ceramide synthesis by Fumonisin B1 ameliorated palmitate-mediated ER stress in MIN6 cells. When coupled with increased C16:0 ceramides levels in islets that lack FIT2, the data suggests that increased ceramides link β-cell FIT2 loss with enhanced ER stress, adding to current knowledge of FIT2 loss leading to ER lipid imbalance and ER dysregulation (23, 24). We present a pathway for how palmitate drives ER stress, with FIT2 stability and LD accumulation taking the center stage in β-cells. The data presented here go beyond the past correlates of ceramide synthase inhibition improving GSIS in palmitate-treated β-cells and reducing apoptosis in TAG-treated macrophages (29, 39). Failure to sequester lipids away from the ER increases substrates for ER-localized enzymes serine palmitoyltransferase and ceramide synthase to enhance ER ceramide levels (40, 41). Ceramide accumulation has been suggested to change the biophysical properties of ER membrane rafts, induce defective ER protein export, disrupt ER Ca^2+^ homeostasis and subsequently trigger ER stress-mediated apoptosis (42, 43). The strongest evidence of lipid-induced ER stress because of FIT2 loss is the observed increased presence of CHOP, a known pre-apoptotic death signal. Thus, under lipotoxic conditions, FIT2 and LDs function critically to sequester and store FAs in their less toxic esterified forms thereby preventing ceramides, especially C16:0, from accumulating within the ER.

In a contrasting situation where LDs were allowed to accumulate in β-cells through desnutrin (a TAG hydrolase) ablation, a similar blunted GSIS response was observed (44). These results seem contradictory to what is reported here as they suggest that the converse, intracellular LD accumulation, is detrimental to β-cells. However, it is noteworthy that the significant driver for the reduced GSIS in the desnutrin knockout mice is reduced fatty acid utilization by mitochondria, rather than enhanced β-cell ER stress. The latter mechanism that supports our findings. To reconcile this perceived difference, it is important to note that while LD formation and lipid sequestration away from the ER mitigates lipotoxicity, this is perhaps on the premise that LD utilization and TAG hydrolysis remains unaffected. Here, further work is required to delineate the dynamics of LD accumulation and LD turnover, especially since higher number of LDs has been observed in human diabetic β-cells (16). Nevertheless, it is increasingly clear that the formation of LDs, especially during lipotoxic conditions, is critical for preventing ER lipid accumulation and ER stress.

A fundamental mechanism governing LD loss is the post-translational modification of FIT2 by palmitate (and stearate) and its subsequent degradation in β-cells. This may explain the lack of FIT2 being mentioned in any GWAS or islet RNA-seq related studies, although a recent diabetes meta-analysis study identified an East Asian diabetes-associated loci (RS6017317) in the regulatory region of FIT2 (45). The molecular switch of this protein, and a potential therapeutic strategy, likely involves protein stability maintenance at the ER, as evidenced by a partial rescue of lipotoxicity through FIT2 overexpression alone. Palmitoylation commonly occurs on transmembrane proteins, affecting protein stability and subcellular localization (46, 47). Interfering with palmitoylation either pharmacologically or through site-directed mutagenesis reduced palmitate-induced FIT2 degradation, and a similar mechanism was reported in other transmembrane proteins such as TBC1D3 and CDCP1 (48, 49). We further identified ERAD-C as the most likely pathway responsible for FIT2 degradation with increased MARCH6 and FIT2 protein association in the presence of palmitate and reduced FIT2 degradation with MARCH6 knockdown. Mutating predicted S-acylated cysteine residues abrogated FIT2 palmitoylation and degradation. Further molecular work correlating degree of FIT2 palmitoylation with its degradation in both physiology and pathology may help improve our understanding on how FIT2 function is fine-tuned at the protein level.

In conclusion, our results show how physiological levels of dietary SFAs, but not USFAs, disable LD biogenesis in β-cells. Palmitoylation of FIT2 leads to its degradation likely through ERAD-C mediated mechanisms, with increased association between FIT2 and MARCH6. Loss of FIT2 leads to a significant reduction in β-cell LDs and increases ceramide accumulation. As a consequence, ER stress hampers β-cell function and increases β-cell apoptosis (i.e. lipotoxicity). Beyond FIT2 and the understanding of how SFAs are more damaging than UFAs to cells, our results show that restoration of LD formation, especially in a lipotoxic milieu such as obesity and diabetes, is of considerable therapeutic value for preventing β-cell dysfunction and loss.

## MATERIALS & METHODS

### Mice

β-cell specific FIT2 knockout mice (β-FIT2KO, KO) were generated using the Cre-lox recombination system. Mice with *floxed* FIT2 (FIT2fl/fl, FL) (21) were bred with mice expressing Cre-recombinase, under control of the rat insulin promoter (B6.Cg-Tg(Ins2-cre)25Mgn/J). Male mice were used in all animal experiments. β-FIT2KO and their corresponding *floxed* littermates (FIT2fl/fl) were fed with western-type (West) diet (D12079B, Research Diets) from 5 weeks of age for a further 25 weeks. B6.BKS (D)-Lepr^db^/J (db/db) mice were compared with their age-matched heterozygous littermates (db/+) at 16 weeks. Mice were housed in a facility with a 12-h light-dark cycle and with food and water available *ad libitum*. All the animal experiments and protocols were approved by the Institutional Animal Care and Use Committee of SingHealth (IACUC SingHealth # 2013/SHS/816) and of NTU Singapore (A0373).

### Islet isolation, dispersion and cell culture

Mouse pancreatic islets were isolated by perfusing the pancreas through the common bile duct with collagenase as previously described (50). The isolated islets were dispersed into single cells by incubation with Accutase^®^ at 37°C for 3-5 min. The primary islet cells were cultured in CMRL medium supplemented with 10% heat-inactivated fetal bovine serum (FBS), 100 U/ml penicillin and 100 μg/ml streptomycin. MIN6 cells (passages 30–35) were cultured in DMEM containing 10% heat-inactivated fetal bovine serum, 25 mM glucose, 4 mM L-glutamine, 100 U/ml penicillin, 100 μg/ml streptomycin and 50 μM 2-mercaptoethanol. Cells were maintained at 37 °C under 5% CO2 and 95% air. Transient transfections of MIN6 were performed using Lipofectamine 2000 (ThermoFisher Scientific) according to the manufacturer’s instructions. FFAs (palmitate, stearate, oleate and palmitoleate) were conjugated with FFA-free BSA at a 2:1 molar ratio and used at a final concentration of 300 μM.

### Protein extraction and immunoblotting assays

Cells and isolated islets were lysed in RIPA buffer supplemented with protease inhibitor cocktail. Proteins were separated by SDS-PAGE and transferred onto nitrocellulose membranes. Blocking was performed at room temperature for 1 h in Tris-buffered saline (TBS) with 5% non-fat milk, followed by incubation with the different primary antibodies (described above) in blocking buffer for either 1 h at room temperature or overnight at 4 °C. After several washes with TBS containing 0.5% Tween 20 (TBST), the membranes were incubated with secondary antibodies of anti-mouse/rabbit IgG/HRP (as appropriate) in TBS with 1% non-fat milk. Following several washes, the protein bands were visualized using enhanced chemiluminescence (Cell Signaling Technology) and quantified using ImageJ.

### RNA extraction and quantitative RT-PCR

Total RNA was prepared from tissues or cells using the NucleoSpin RNA II kit (Macherey-Nagel), or prepared from the isolated islets using the NucleoSpin RNA XS kit (Macherey-Nagel). 1 μg of RNA was reversely transcribed using High Capacity cDNA Reverse Transcription Kit (Applied Biosystems) according to the manufacturer’s instructions. Real-time PCR was performed on QuantStudio 6 Flex Real-Time PCR System (Applied Biosystems) using Power SYBR Green PCR Master Mix (Applied Biosystems). The PCR primers used are summarized in Supplementary Table 3.

### RNA-seq library preparation and data processing

Pancreatic islets were isolated from 12-week old wild type (WT) and β-cell specific FIT2 knockout (β-FIT2KO, KO) mice in quintuplicate. Isolated pancreatic islets then cultured in complete CMRL medium overnight for recovery. Total RNA was harvested using RNeasy Plus Mini Kit (Qiagen) followed by RNA-seq library construction. Genes were considered to be significantly differentially expressed when false discovery rate (FDR) ≤ 0.05, with FPKM ≥ 1 in one sample group retained for subsequent analysis. GO clustering enrichment analysis was carried out on differential genes using the Functional Annotation tool in DAVID version 6.8 (51, 52) under medium stringency for all default annotation categories except protein domains. We identified significant clustered groups having group enrichment scores of ≥ 1.3, with higher scores indicative of more significant annotated GO clusters. Heatmaps were generated with R package pheatmap version 1.0.10 (https://cran.r-project.org/web/packages/pheatmap/index.html).

### Immunofluorescence

Cryosections of 10 µm thickness from fixed and cryopreserved pancreas were used for immunofluorescence analysis. Sections were rinsed with TBS, permeabilized, and blocked with 10% normal goat serum plus 0.2% Triton X-100 in TBS for 1 h at room temperature and then incubated overnight with primary antibodies at 4°C in a humidified atmosphere. After gentle washing with TBS and incubating with fluorescence secondary antibodies for 1 h at room temperature, sections were mounted with Vecta Mount solution (Vector Labs) and multiple Z-stack images were obtained using confocal imaging (Leica) and subsequently quantified using Image-J). Briefly, the average fluorescence intensity of the staining, within insulin-positive ROI, was quantified, corrected by subtraction of average background intensity, and normalized to its control. For LD staining, cryosections were stained with BODIPY 493/503 (0.01mg/ml) and DAPI (5 μg/ml) for 15 min at RT. For propidium iodide (PI) staining, cells were incubated with 10 μg/ml PI and 5 μg/ml DAPI in medium for 1h at 37 °C followed by RT fixation. Images were captured using a SP8 confocal microscope (Leica). BODIPY-positive puncta per cell and percentage of PI-positive cells were quantified using Image-J Prism 7 (GraphPad).

### Total islet and pancreatic insulin content

To determine islet insulin content, 10 isolated islets were washed twice with ice-cold D-PBS and then lysed with RIPA buffer. Insulin and protein content of the lysate were measured using Mouse Insulin ELISA (Mercodia) and BCA assay (ThermoFisher Scientific), respectively. Total islet insulin content was normalized to total protein content. To determine pancreatic insulin content, half of the whole pancreas was collected and placed into 5 ml Acid-Ethanol solution (1.5% HCl in 70% EtOH) overnight at −20°C. Tissue was then homogenized and incubated in the same solution overnight at −20°C. Supernatant was collected by centrifugation at 3000 *x g* for 10 min at 4°C, followed by neutralization with 1 M Tris pH 7.5 at 1:1 (*vol/vol*). Insulin and protein content of the neutralized solution were measured. Total pancreatic insulin content was normalized by total protein content.

### Glucose Stimulated Insulin Secretion (GSIS)

MIN6 cells were pre-incubated at 37°C for 2 h in Krebs Ringer Bicarbonate (KRBH) buffer (119 mM NaCl, 4.74 mM KCl, 2.54 mM CaCl_2_, 1.19 mM MgCl_2_, 1.19 mM KH_2_PO_4_, 25 mM NaHCO_3_, and 10 mM HEPES, 0.1% BSA, 0.1 mM glucose, pH 7.4). At experiment start, fresh KRBH was added for 30 min, followed by 25 mM glucose KRBH at 37°C for another 30 min. The supernatant was collected for insulin measurements and normalized to total protein. Isolated islets were 2 h pre-incubated with buffer (125 mM NaCl, 5.9 mM KCl, 2.56 mM CaCl_2_, 1.2 mM MgCl_2_, 25 mM HEPES, 0.1% BSA and 2.8 mM glucose, pH 7.4) at 37°C. 50 islets were placed between two layers of bio-gel (Bio-Rad) in a perifusion chamber and perifused at a flow rate of 200 μL/min, at 37°C, for 20 min prior to the start of sample collection. Islets were perifused for 10 min with 2.8 mM glucose buffer, followed by 10 min with 16.7 mM glucose buffer. At the end of the perifusion, islets were lysed in RIPA buffer and total insulin content was measured for normalization. For *in vivo* GSIS, mice were starved for 6 h and blood samples were collected from tail vein to determine basal insulin levels. Mice were then injected with 3 g glucose/kg of body weight, intraperitoneally. Blood samples were collected from the tail vein 15 min after glucose injection. Total blood was centrifuged at 10,000 *x g* for 10 min and the supernatant (serum) was collected for insulin measurements using Mouse Insulin ELISA (Mercodia).

### Insulin and glucose tolerance tests

Prior to GTT or ITT, mice were fasted for 6 h with *ad libitum* access to water and then administered glucose (2 g/kg) or insulin (0.75 unit/kg) intraperitoneally. Blood glucose was measured in blood samples collected from tail vein using a glucometer (Accu-Chek Performa Nano System).

### Ceramides analysis

Pancreatic islets were resuspended in 100 μL of butanol/methanol (1:1, v:v) spiked with 66.07 nM of C8 Ceramide (d18:1/8:0) purchased from Avanti Polar Lipids (860508). The cells were sonicated for 30 min followed by centrifugation at 14,000 g for 10 min at 22 °C. The supernatant was transferred to MS vials for analysis. The samples were analysed using an Agilent 1290 series UHPLC system connected to an Agilent 6495 QQQ mass spectrometer after separation on a ZORBAX Eclipse plus C18 column (2.1 x 50 mm, 1.8 µm, 95 Å, Agilent) at 40 °C. The injection volume was 2 µL. Solvent A consisted of 60% water/40% acetonitrile (v/v) with 10mM ammonium formate; solvent B consisted of 90% isopropanol/10% acetonitrile (v/v) with 10mM ammonium formate. The gradient started with a flow rate of 0.4 mL/min at 20%B and increased to 60% B at 2 min, 100% B at 7 min, held at 100% B until 9 min, followed by equilibration with 20% B from 9.01 min until 10.8 min. The column effluent was introduced to the Agilent 6495 QQQ mass spectrometer via AJS-ESI ion source operating under the following conditions: Gas temperature, 200°C; gas flow, 14 L/min; nebulizer, 20 psi; sheath gas temperature, 250°C; sheath gas flow, 11 L/min; capillary, 3500 V. Mass spectrometry analysis was performed in positive ion mode with dynamic scheduled multiple reaction monitoring (dMRM). Mass spectrometry settings, LC-MS gradient and MRM transitions for each lipid class were adapted from a previously published method (53). Data analysis was performed on Agilent MassHunter Quantitative analysis software. Relative quantitation was based on one-point calibration with Ceramide d18:1/8:0. The data were further normalized to the total number of islets in each sample.

### Co-immunoprecipitation assay for FIT2-MARCH6 interaction

MIN6 cells, transiently co-transfected with pcDNA3.1-FIT2/V5-His and GFP-tagged MARCH6 (BC059190) Mouse Tagged ORF Clone (Origene), were pre-treated with 10 µM MG132 (Cell Signaling Technology) for 2 h and subsequently treated with 300 µM palmitate (BSA-conjugated) for 6 h. Cells were then lysed in IP lysis buffer (ThermoFisher Scientific), supplemented with protease inhibitor cocktail. Co-immunoprecipitation assay was then performed on the protein lysates using Pierce™ Classic IP Kit (ThermoFisher Scientific) with either Anti-6X His tag antibody (Abcam) or Rabbit IgG (Cell Signaling Technology) in accordance with the manufacturer’s instructions. The eluted fraction was then subjected to immunoblotting assay and MARCH6 was detected using Mouse monoclonal turboGFP antibody (Origene).

### S-Palmitoylation assay

MIN6 cells, transiently transfected with pcDNA3.1-FIT2/V5-His, were pre-treated with 10 µM MG132 (Cell Signaling Technology) for 2 h and thereafter treated with 300 µM palmitate (BSA-conjugated) for 6 h. S-palmitoylation assay was then performed using CAPTUREome™ S-Palmitoylated Protein Kit (Badrilla) in accordance with manufacturer’s instruction. Briefly, cell lysis and free thiol blocking (blocking of free thiols on non-palmitoylated cysteine residue) was performed by incubating cells with the provided lysis buffer for 4 h with constant shaking. Proteins were then precipitated using acetone, and subsequently resuspended in the provided binding buffer. 30 µl of resuspended proteins were collected as the Input Fraction (IF), while the remainder was split into two tubes: “experimental” and “negative control”. In the experimental tube, proteins were treated with thioester cleavage reagent, which cleaves off acyl groups from the protein, resulting in the exposure of a free thiol group. Treated samples were then subjected to CAPTUREome™ resin for 2.5 h at room temperature, where proteins with free thiols were captured by the resin. 50 µl of flowthroughs were collected as the cleaved unbound fraction (cUF). S-acylated proteins, which were captured by the CAPTUREome™ resin, were eluted by incubating the resin in 50 µl of 2x laemmli buffer at 60°C for 10 min (cleaved Bound Fraction (cBF)). In the negative control tube, proteins were treated with acyl-preservation reagent, which preserves acyl groups on the protein. Treated samples were then subjected to CAPTUREome™ resin for 2.5 h at room temperature. 50 µl of flowthroughs were collected as preserved unbound fraction (pUF). Proteins were eluted from the CAPTUREome™ resin by incubation of resin with 50 µl of 2x laemmli buffer at 60°C for 10 min (preserved Bound Fraction (pBF)).

### Statistical analysis

Data tested for normality (Shapiro-Wilk test) are expressed as mean ± SEM. Parametric analysis (using Student’s t-test) was used to determine statistical difference and P values < 0.05 were considered as statistically significant (Prism 7, GraphPad).

## DATA AVAILABILITY

RNA sequencing data reported in this paper is available at NCBI GEO Accession no.: GSE133939. All data mentioned in this paper will be placed on a data repository. Authors declare no primary datasets and computer codes linked to this study.

## ACKNOWLEDGMENTS

The mouse pancreatic β-cell line MIN6 was kindly provided by Dr. Jun-ichi Miyazaki, Osaka University, Japan. This work was supported by the Singapore Ministry of Education under its Singapore Ministry of Education Academic Research Fund Tier 2 and Tier 1 (MOE2015-T2-2-087 and 2017-T1-001-220, 2019-T1-001-059) (Y.A.) and the Lee Kong Chian School of Medicine, Nanyang Technological University Singapore Start-up Grant (to Y.A. and P.O.B.) This work is also partly supported by the LKCMedicine Healthcare Research Fund (Diabetes Research), established through the generous support of alumni of Nanyang Technological University, Singapore. C.H. is supported by the Lee Kong Chian School of Medicine and the Nanyang President’s Graduate Scholarship, Nanyang Technological University, Singapore. J.N.F. is supported by the Singapore National Research Foundation Fellowship (NRF-NRFF2016-03). Work in M.R.W. and F.T. laboratory is supported by grants from the National University of Singapore via the Life Sciences Institute (LSI), the National Research Foundation (NRF, NRFI2015-05 and NRFSBP-P4) and the NRF and A*STAR IAF-ICP I1901E0040. P.O.B. is additionally supported by the Swedish Research Council, the Family Erling-Persson Foundation, the Novo Nordisk Foundation, the Stichting af Jochnick Foundation, the Swedish Diabetes Association, the Scandia Insurance Company Ltd., Diabetes Research and Wellness Foundation, Berth von Kantzow’s Foundation, the Strategic Research Program in Diabetes at Karolinska Institutet, the ERC-“Advanced Grant” (EYELETS), and the Center of Excellence-International Collaboration Initiative Grant (China). X.Z. is currently supported by the National Natural Science Foundation of China (82070846). The authors would also like to thank the A*STAR Microscopy Platform (Singapore) for assistance in sample processing and for the electron microscopy analysis in this study.

## CONTRIBUTIONS

X.Z. and Y.A. designed the study and wrote the manuscript. D.S. and P.O.B. provided input on floxed-FIT2 mice studies. X.Z, C.H, M.C., S.M. and O.S. performed the experiments. E.G.C., M.M.L. and J.N.F. analysed the RNA-sequencing data. C.H., D.S., J.N.F., P.O.B., F.T., M.R.W. and Y.A. edited the manuscript.

---

**Supplementary Fig 1:**
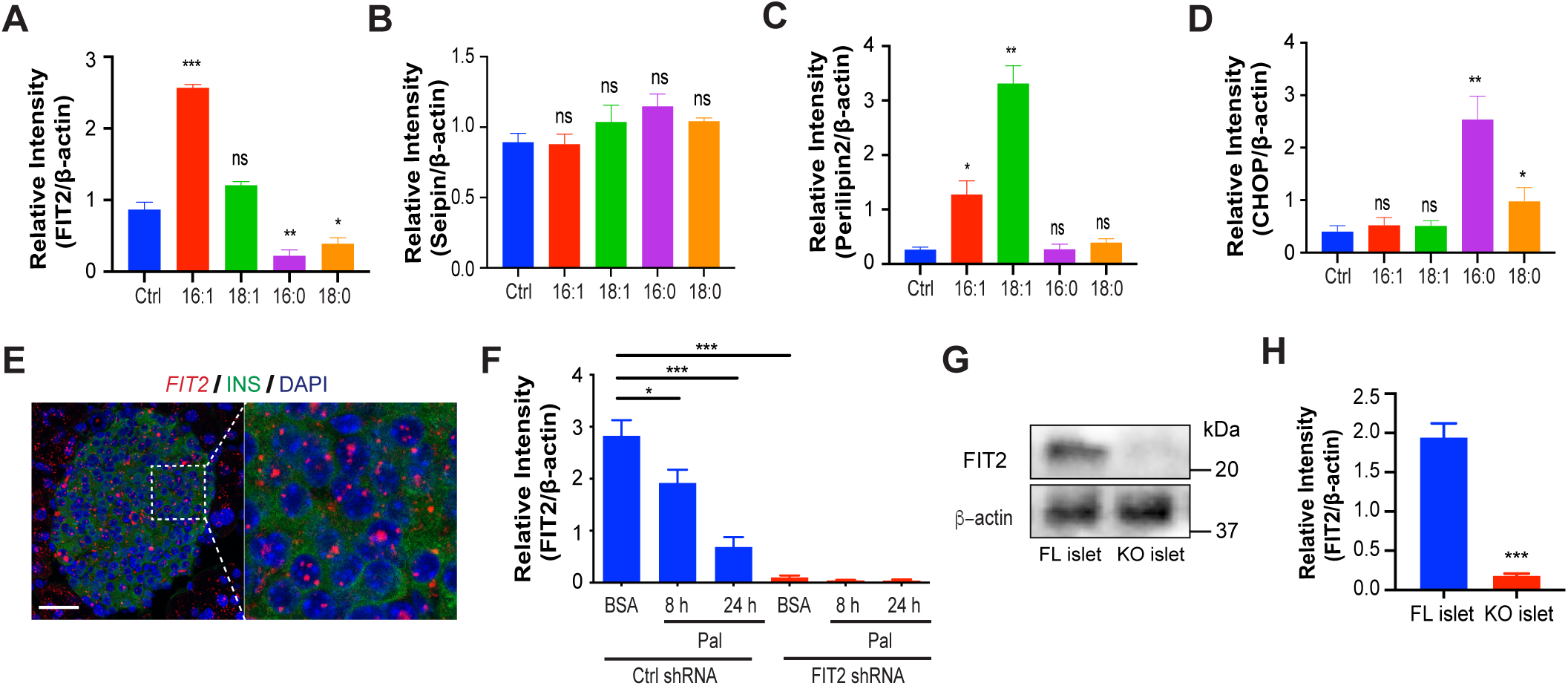
Relative levels of LD formation proteins following FA treatment as well as analysis of FIT2 in islets of βFIT2KO mice. **A-D**, Semi-quantitation of immunoblots for FIT2, Seipin, Perilipin2 and CHOP protein levels in MIN6 cells treated with palmitoleate (16:1), oleate (18:1), palmitate (16:0), stearate (18:0) (300 µM, 24 h) or BSA control (N=3-4). **E**, Representative i*n situ* hybridization image for *FIT2* mRNA (red) combined with immunohistochemistry for insulin (green) in pancreatic cryosections from C57BL/6 mice. DAPI was used to counterstain the cell nucleus (blue). Expanded image (right) of the demarcated section. Images shown are maximum-intensity projections. Scale bar = 50 µm. **F,** Semi-quantitation of immunoblot analysis of FIT2 expression in vector only (Ctrl shRNA) or stable FIT2 knockdown (FIT2 shRNA) MIN6 cells treated with or without palmitate (300 µM). **G**, Representative immunoblot and **H**, corresponding semi-quantitation of FIT2 protein in isolated islets from floxed control (FL) and βFIT2KO (KO) mice (12 wk-old male, N=7). Values shown are mean ± SEM; ns, not significant; *, P < 0.05; **, P < 0.01; ***, P < 0.001 relative to control (two-tailed Student’s t-test).

**Supplementary Fig 2:**
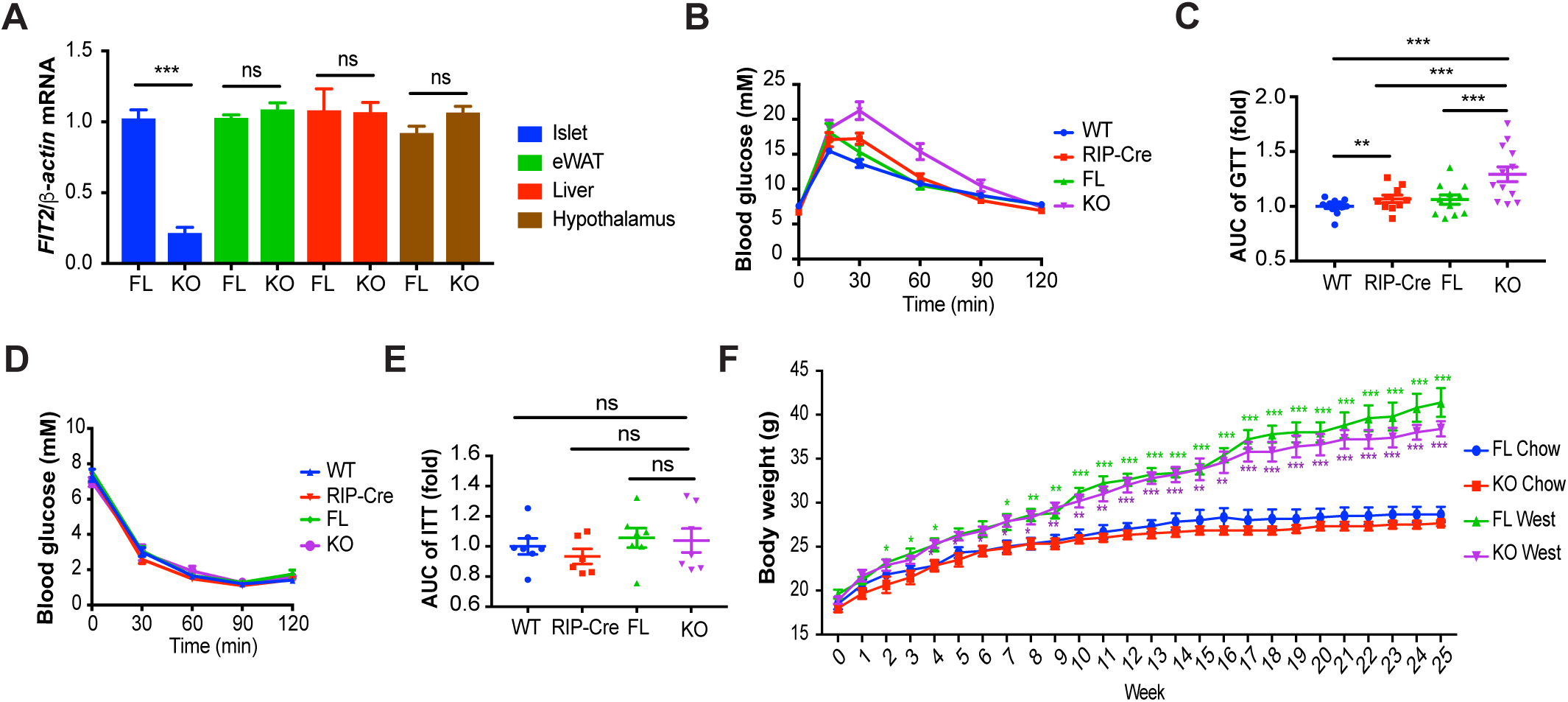
Extended phenotype description of βFIT2KO mice. **A**, Gene expression (qPCR) analysis of *FIT2* mRNA levels in pancreatic islets, adipose tissue, liver and hypothalamus of floxed control (FL) and βFIT2KO (KO) mice (12 wk-old male, N=3-4). **B**, IPGTT on FL, KO, Rip-Cre and WT mice and **C**, corresponding area under the curve (AUC) of blood glucose levels (12-16 wk-old male, N=10-13). **D**, ITT on FL, KO, Rip-Cre and WT mice and **E**, corresponding AUC of blood glucose levels (12-16 wk-old male, N=7). **H**, Body weight of floxed control (FL) and βFIT2KO (KO) fed with chow diet (FL Chow, KO Chow) or West diet (FL West, KO West) for 25 weeks (N=6). Values shown are mean ± SEM; *, P < 0.05; **, P < 0.01; ***, P < 0.001 relative to control (two-tailed Student’s t-test).

**Supplementary Fig 3:**
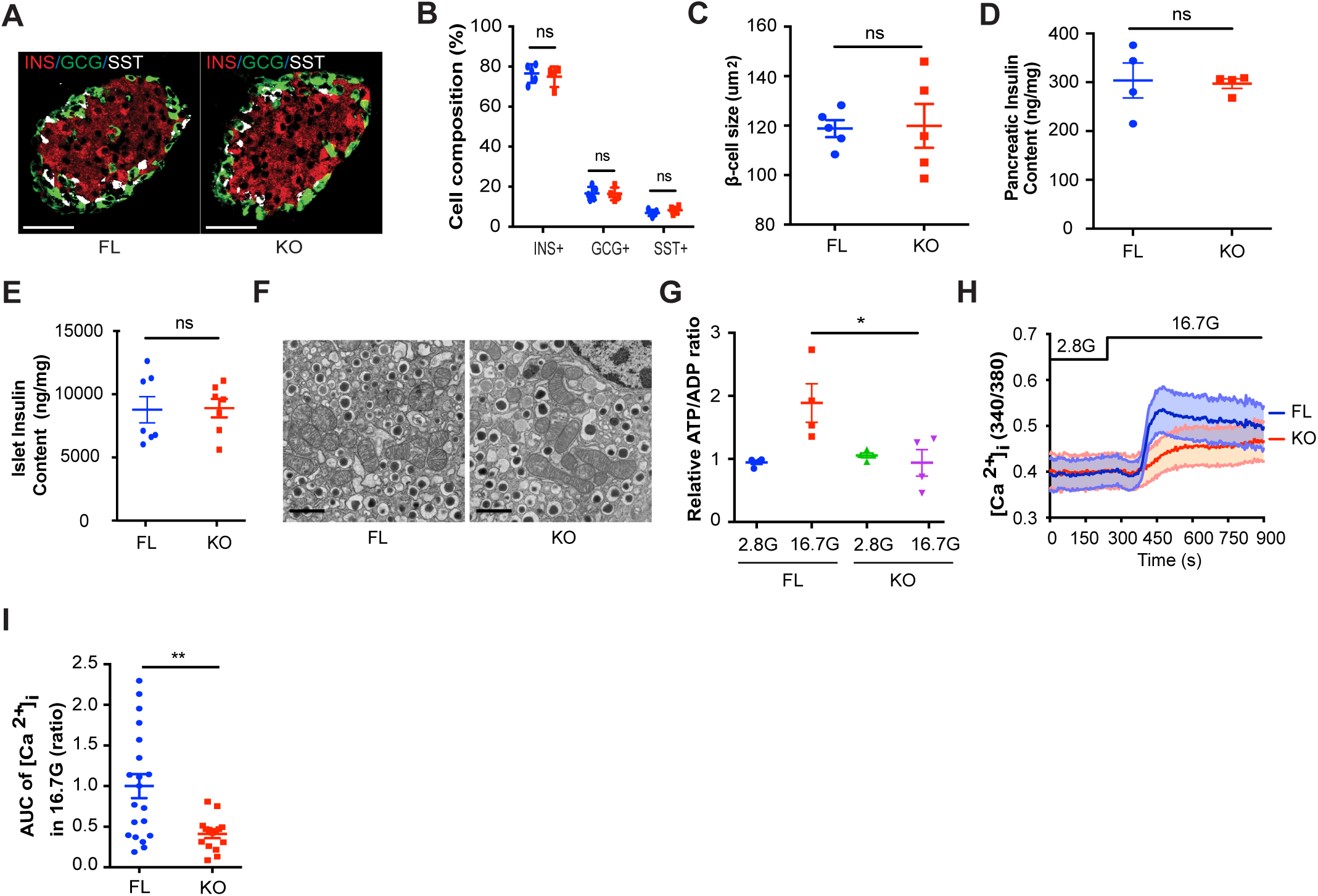
Islet characteristics of βFIT2KO mice. **A**, Representative immunostaining for insulin (red), glucagon (green) and somatostatin (white) in pancreas sections from FL and KO mice (12 wk-old male). Scale bar = 50 µm. **B, C,** Cell-type distribution and β-cell size analysis of *in situ* islets in pancreas sections from floxed control (FL) and βFIT2KO (KO) mice (12 wk-old male, N=4-5). **D, E**, Measurements of total pancreas (N=4) and islet (N=7) insulin content. **F**, Representative Transmission Electron Microscope (TEM) images of insulin granules of FL and KO pancreata (N=3). Scale bar = 1 µm. **G**, ATP/ADP ratio of FL and KO islets in 2.8 mM glucose (2.8 G) followed by stimulation with 16.7 mM glucose (16.7 G). (N=4). **H**, Measurements of changes in [Ca^2+^]_i_ in FL and KO islets exposed to varying concentrations of glucose “G” as indicated (15-20 islets from four independent experiments were analyzed per group). **I**, AUC of [Ca^2+^]_i_ at 16.7 G (n=15-20). Values shown are mean ± SEM; ns, not significant; *, P < 0.05; **, P < 0.01; ***, P < 0.001 relative to control (two-tailed Student’s t-test).

**Supplementary Fig 4:**
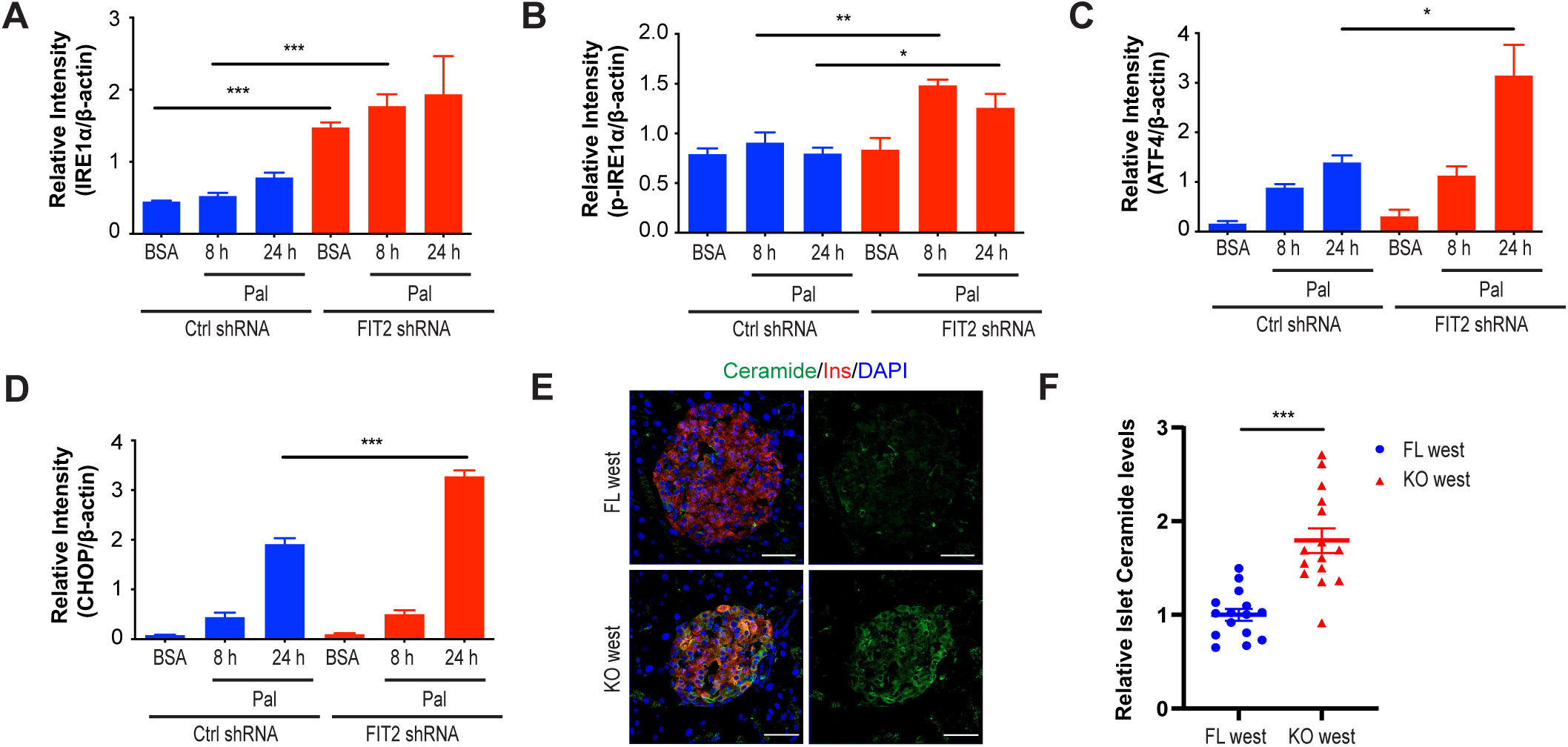
Relative levels of ER stress proteins and ceramide immunodetection in pancreas of βFIT2KO mice. **A-D**, Semi-quantitation of immunoblot analysis of IRE1α, p-IRE1α, ATF4 and CHOP protein levels in scrambled (Ctrl shRNA) or FIT2 knockdown (FIT2 shRNA) MIN6 cells in the absence (BSA) or presence of palmitate (300 µM). **E,** Representative immunostaining image for insulin (red), ceramide (green) and DAPI (blue) in pancreas sections from 25 wk of western diet fed FL and KO mice. Scale bar = 50 µm. **F,** Quantitation of ceramide fluorescence intensities from floxed control (FL) and βFIT2KO (KO) islets of 25 wk of western diet fed mice (normalised to FL) using Image-J. Values shown are mean ± SEM; ns, not significant; *, P < 0.05; **, P < 0.01; ***, P < 0.001 relative to control (two-tailed Student’s t-test).

**Supplementary Fig 5:**
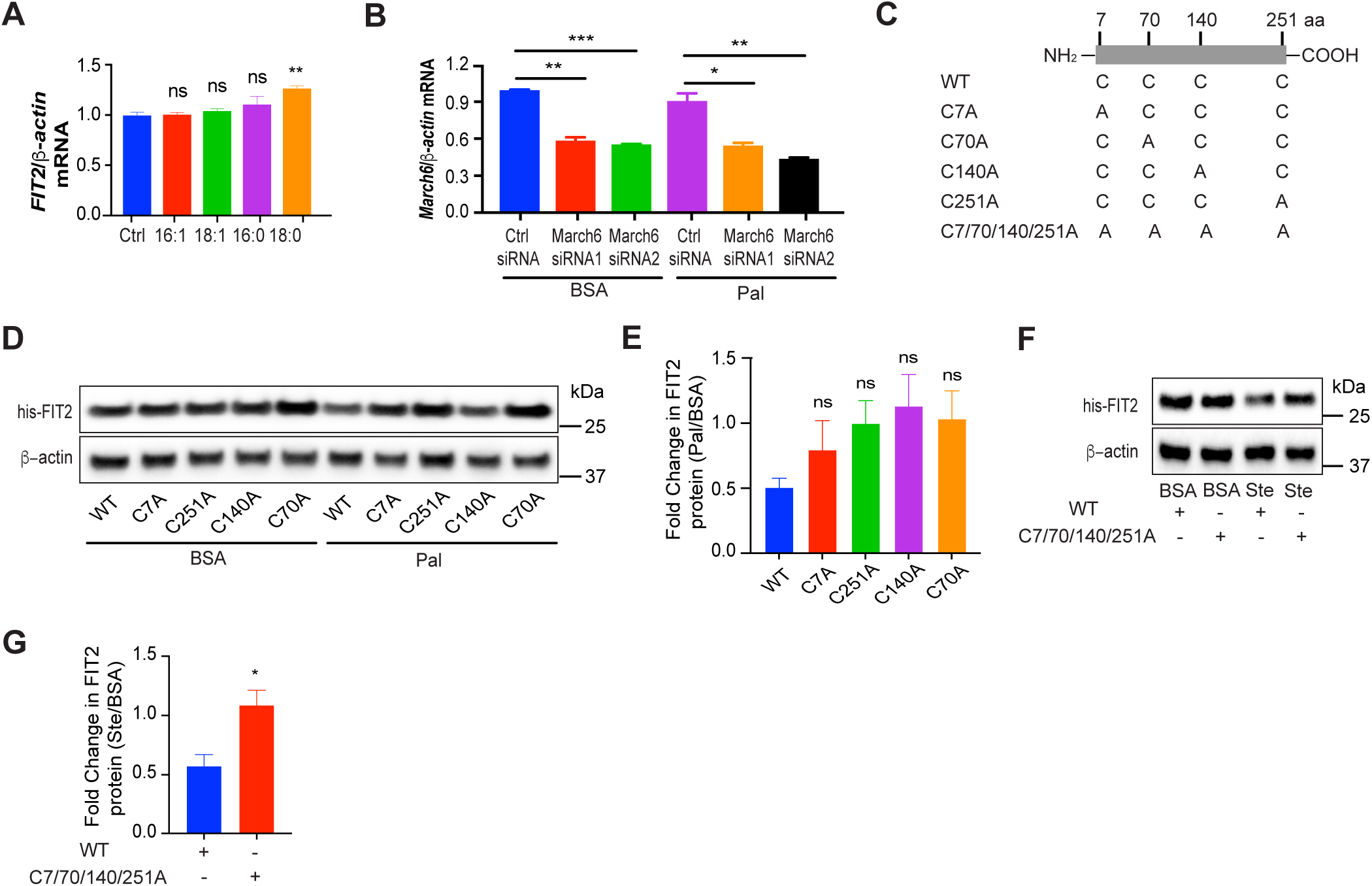
Relative changes in gene and protein levels of FIT2 following exposure to different FAs and separately on *March6* knockdown efficiency in MIN6 cells. **A**, Relative *FIT2* mRNA levels (qPCR) in MIN6 cells treated with BSA or different FAs (300 µM) for 24 h (N=3). **B**, MIN6 cells were transiently transfected with a scrambled (ctrl siRNA) or different siRNA oligonucleotides targeting *March6*. 36 h after transfection, cells were treated with either BSA or palmitate (300 µM) for 4 h. **B**, Relative *March6* gene expression levels following siRNA mediated knockdown. **C**, Schematic representation of point mutants (C → A) of FIT2. **D-G**, Representative immunoblot analysis and semi-quantification of FIT2 protein levels in MIN6 cells transfected with pcDNA3.1-FIT2/V5-His (WT) or mutant followed by treatment with BSA, palmitate (300 µM) or stearate (300 µM) for 24 h (N=3-4). Values shown are mean ± SEM; ns, not significant; *, P < 0.05; **, P < 0.01; ***, P < 0.001 relative to control (two-tailed Student’s t-test).

